# Inhibitory control performance is repeatable across years and contexts in a wild bird population

**DOI:** 10.1101/2021.07.15.452478

**Authors:** Gabrielle L. Davidson, Michael S. Reichert, Jennifer R. Coomes, Ipek G. Kulahci, Iván de la Hera, John L. Quinn

## Abstract

Inhibitory control is one of several cognitive mechanisms required for self-regulation, decision making and attention towards tasks. Linked to a variety of maladaptive behaviours in humans, inhibitory control is expected to influence behavioural plasticity in animals in the context of foraging, social interaction, or responses to sudden changes in the environment. One widely used cognitive assay, the ‘detour task’, putatively tests inhibitory control. In this task, subjects must avoid impulsively touching transparent barriers positioned in front of food, and instead access the food by an alternative but known route. Recently it has been suggested that the detour task is unreliable and measures factors unrelated to inhibitory control, including motivation, previous experience and persistence. Consequently, there is growing uncertainty as to whether this task leads to erroneous interpretations about animal cognition and its links with socio-ecological traits. To address these outstanding concerns, we designed a variant of the detour task for wild great tits (*Parus major*) and deployed it at the nesting site of the same individuals across two spring seasons. This approach eliminated the use of food rewards, limited social confounds, and maximised motivation. We compared task performance in the wild with their performance in captivity when tested using the classical cylinder detour task during the non-breeding season. Task performance was temporally and contextually repeatable, and none of the confounds had any significant effect on performance, nor did they drive any of the observed repeatable differences among individuals. These results support the hypothesis that our assays captured intrinsic differences in inhibitory control. Instead of ‘throwing the detour task out with the bathwater’, we suggest confounds are likely system and experimental-design specific, and that assays for this potentially fundamental but largely overlooked source of behavioural plasticity in animal populations, should be validated and refined for each study system.

## Introduction

Inhibitory control is a well-known form of self-regulation that affects decision making and is linked to a variety of maladaptive behaviours in humans, including addiction (Jentsch & Taylor, 1999; Loeber & Duka, 2009), obesity (Epstein et al., 2008; Houben et al., 2014) and asocial emotional responses (Tang & Schmeichel, 2014). Inhibitory control is increasingly being used to study animal behaviour due to its potential role in socio-ecological processes (reviewed in Kabadayi et al., 2018), including foraging plasticity (Coomes et al., 2020), dietary breadth (MacLean et al., 2014; Van Horik et al., 2018), and social interactions (Amici et al., 2013, 2018; Reddy et al., 2015). Moreover, inhibitory control is likely responsive to selection given its links with brain size in primates, and its heritability in humans and birds (Friedman et al., 2008; Langley et al., 2020). However, the inferential power of these studies is dependent on cognitive assays that accurately characterise inhibitory control and whether these assays are applicable to how animals behave in natural settings. Recently, researchers have taken a variety of classical cognitive tasks to the field (e.g. Johnson-Ulrich et al., 2020; Morand-Ferron et al., 2011, 2015, 2016; Muth et al., 2018; Reichert et al., 2020; Sonnenberg et al., 2019; Toledo et al., 2020) where cognition can be assayed under natural conditions with high ecological validity, and welfare concerns and administrative burdens associated with housing animals in captivity are reduced. However, inhibitory control assays in the wild are scant, and to date rely on experimenters in close proximity to habituated animals (Ashton et al., 2018; Shaw et al., 2015). Moreover, there remains uncertainty regarding what is being measured in inhibitory control tasks in animals. For example, variation in performance on one assay of inhibitory control does not always predict performance in another. Possible reasons for this include that inhibitory control is a composite trait, and variations of the task can measure different components of inhibitory control (Bari & Robbins, 2013; Völter et al., 2018); that inhibitory control is tightly linked with many other decision-making processes, such as attention and task switching (Bari & Robbins, 2013); and because non-cognitive processes (e.g. previous experience) confound task performance (Jelbert et al., 2016; van Horik et al., 2020; Van Horik et al., 2018). More generally, the extent to which extraneous variables contribute to or confound cognitive performance may affect captive-bred, temporarily-captive wild individuals, and free-living animals differentially (Morand-Ferron et al., 2016), but comparisons across such experimental contexts have rarely been examined empirically for any cognitive trait (see Benson-Amram et al., 2013; Cauchoix et al., 2017; Forss et al., 2015; McCune et al., 2019; Morand-Ferron et al., 2011; Mouchet & Dingemanse, 2021 for some examples).

One way of measuring inhibitory control is the widely used ‘detour’ task. In this task, subjects must avoid and move around a transparent barrier to retrieve a food reward that is positioned directly in front of them, but behind the barrier (Diamond, 1990). A central premise of this task is that the visible reward generates a strong prepotent impulse to approach it directly. This impulse must be inhibited, and the subject must instead move in a direction away from the reward in order to detour around the barrier successfully. A practical advantage of the detour task is that it is less laborious than other cognitive assays, because animals quickly pass the habituation and training phases, and the test phases can be achieved within 10 trials (MacLean et al., 2014) or even three trials (Van Horik et al., 2018). Therefore the detour task has the potential to be a valuable field paradigm for addressing key questions regarding the causes and consequences of individual differences in cognition, provided it is a reliable measure of inhibitory control (Shaw & Schmelz, 2017; Thornton et al., 2014). However, because the classic detour task is dependent on an experimenter re-baiting the apparatus with a food reward, its scope for field-based assays has thus far been restricted to wild, human-habituated New Zealand robins and Australian magpies (e.g. Ashton et al., 2018; Shaw, 2017b). Another limitation for cognitive tasks in the field is that invariably these tasks involve providing food rewards where many species visit in groups. Consequently, individual performance is likely subject to social interference, for example through social learning or competition from other group members (Cole & Quinn, 2012; Morand-Ferron & Quinn, 2011; Reichert et al., 2020). To address social interference, individuals can be temporarily brought into captivity, where they are housed in relative isolation. While this approach may solve confounds linked to sociality, they could equally generate different confounding effects on performance, for example through stress and motivation (Butler et al., 2006).

Cognition is challenging to assay because the expression of cognitive capabilities is potentially always confounded by some other factor, such as motivation, personality and persistence (Morand-Ferron et al., 2016). This is likely also true for inhibitory control, which plays a role in executive functioning and decision making, works in tandem with additional executive functions such as attention towards the environment (Bari & Robbins, 2013), and belongs to a domain-general set of brain networks involving multiple interacting cognitive processes (Hampshire & Sharp, 2015). The extent to which cognitive tasks that target inhibitory control tap into additional cognitive and non-cognitive processes may therefore be dependent on the type of task used (e.g. Völter et al., 2018), the context (e.g. captivity vs the wild), and temporal aspects (e.g. season). One approach used for quantifying putative cognitive traits generally has been to control for confounds experimentally (e.g. through food deprivation and training), or to independently measure multiple behavioural traits and test their effect on cognitive performance statistically (e.g. Cooke et al., 2021; Reichert et al., 2020). In wild populations, pinpointing confounds is a critical step towards accurately assaying inhibitory control, and interpreting links with functional behaviour, life-history, and evolution.

Examining the consistency of task performance has the potential to shed light on task validity. Typically consistency is quantified by estimating repeatability, that is, by estimating the proportion of variance in a trait measured multiple times that explains between-individual, rather than within-individual, variation within the population. Significant repeatability suggests intrinsic among individual differences in the behaviour, and although it sets the upper limit of heritability (Dohm, 2002), equally those individual differences could be caused by permanent environment effects (Wilson, 2018). Repeatability may not necessarily be strong proof of task validity since the task could be consistently measuring the same confounds, especially when using the same task under the same conditions over time. For example, problem solving performance was repeatable in great tits but this repeatability was largely explained by experimentally manipulated motivational effects that were present in both repeat measures (Cooke et al., 2021). This is less likely to be a problem, however, when repeat measures are taken under very different conditions (contextual repeatability) that are unlikely to have shared environmental confounds, and especially when intrinsic confounds are also likely to differ (Nakagawa & Schielzeth, 2010). Finally, as discussed above, confounding variables can explain variation in single measures of cognitive performance, but can also act on the between- and, or the within-individual variance component when dealing with multiple measures from the same individual. It follows that a further important test is to establish whether the consistent between-individual differences specifically are driven by these confounding effects (Cooke et al., 2021; Nakagawa & Schielzeth, 2010). Demonstrating repeatability (temporal) and contextual repeatability in performance on the detour task would provide compelling support for a common cognitive basis of these tasks, especially when controlling for potential confounds.

The great tit (*Parus major*) provides a model species in ecology and evolution field studies, and has also been used for exploring the evolutionary ecology of cognitive variation. Great tits breed in nest boxes and can be fitted with Passive Integrative Transponder (PIT) identification tags for remote detection at nest boxes using Radio Frequency Identification (RFID) technology. They readily engage with experimental apparatuses in the wild and in captivity (Aplin et al., 2015; Cauchard et al., 2013; Cole et al., 2012; Morand-Ferron et al., 2011; Troisi et al., 2020), allowing for comparisons of cognitive performance across different settings. We designed and presented a modified version of the detour task at the nest box over two years during the breeding season involving minimal experimental disturbance, social interference from conspecifics, and no need for food rewards. We also ran a classic version of the task by testing wild-caught great tits in captivity during one winter season. This approach allowed us to test for both contextual and temporal repeatability, and to examine statistically whether environmental or state variables caused between individual differences in detour task performance, rather than the hypothesised cognitive mechanisms underpinning inhibitory control. These variables included experience, persistence, motivation, body size, and habitat. Moreover, because personality has been shown to affect how animals engage and perform on cognitive tasks (Dougherty & Guillette, 2018; Guillette et al., 2017; Sih & Del Giudice, 2012), we also examined whether ‘exploration behaviour in a novel environment’, a commonly used assay of the fast-slow exploration personality-axis (Dingemanse et al., 2002), explained performance on the captive detour task. If these variables did not explain task performance in general, and the repeatability of performance in particular, this would lend support for the detour task’s utility as a robust measure of inhibitory control, where individual performance is not sensitive to bias from extraneous influences.

## Methods

### Field study sites

Our study took place across ten distinct woodland sites in the River Bandon Valley, Co. Cork, Ireland. Five sites were mixed deciduous and five were conifer plantations (Table S1, supplementary materials). Nest boxes, hung at approximately 1.5 metres from the ground, were distributed across these sites at a density of two nest boxes per hectare. Nest boxes were monitored for breeding data (lay dates, incubation period, hatching, clutch size, brood size and number of fledglings). Adults were trapped at the nest box between day 10 and 13 post-hatching to measure biometrics (including weight, wing and tarsus length), and to tag individuals with a PIT tag and a coloured ring with a unique alpha-numerical code for identification. Chicks were ringed and weighed on day 15 posthatching. Experiments took place between 25^th^ May and 25^th^ June 2017 (hereafter year one), and between 26^th^ May and 15^th^ June 2018 (hereafter year two).

### Aviary

Birds were caught from nine of the Bandon Valley sites described above between January and early March 2018, four of which were from mixed deciduous and five were conifer plantations. Birds were transported from the field sites to the aviary in cloth bags within 2 hours of being caught and brought into captivity for approximately 3 weeks before being released at the site of capture. Birds were housed individually in 60L × 50W × 62H cm enclosures with two perches above the floor of the cage. Each box was individually lit with an LED light bulb on a 10L/14D cycle, and equipped with its own ventilation system. A small viewing window (14cm diameter) at the top half of the box, and an access flap at the base (10cm) allowed for experimenters to watch the birds without being seen, and to change experimental apparatuses with minimal disturbance to the birds. Birds could also be observed on a video monitor connected to cameras fixed to the top of each box. Birds were provided with vitamin-fortified water, sunflower seeds and peanuts ad libitum. Mealworms (*Tenebrio molitor* larvae) were provided at 3 time points each day (morning, afternoon, evening). Wax worms (*Galleria mellonella* larvae) were only provided as experimental rewards.

### Experimental procedures: Detour task in the wild

To measure individuals’ detour task performance in the wild, we designed a detour task for the nest box requiring birds to initially avoid opaque, and subsequently transparent, flexible barriers to gain access to their nest hole. To do so, birds were required to alter their normal flight path (i.e. level with the nest hole), such that they could enter from the gap under the barrier (Figure 1). The experiment consisted of three phases: 1) habituation to the opaque experimental apparatuses, 2) training phase to go under an opaque barrier, and 3) testing phase with a transparent but otherwise similar barrier. In both years, a Perspex cover (20cm×20cm) was positioned horizontally on top of the box during the training and test phases, to provide birds with experience with transparent plastic (cf. Isaksson et al., 2018; Van Horik et al., 2018) and to act as a cover for the transparent barrier in case of rain (although most experiments occurred when there was no rain).

**Figure 1.**
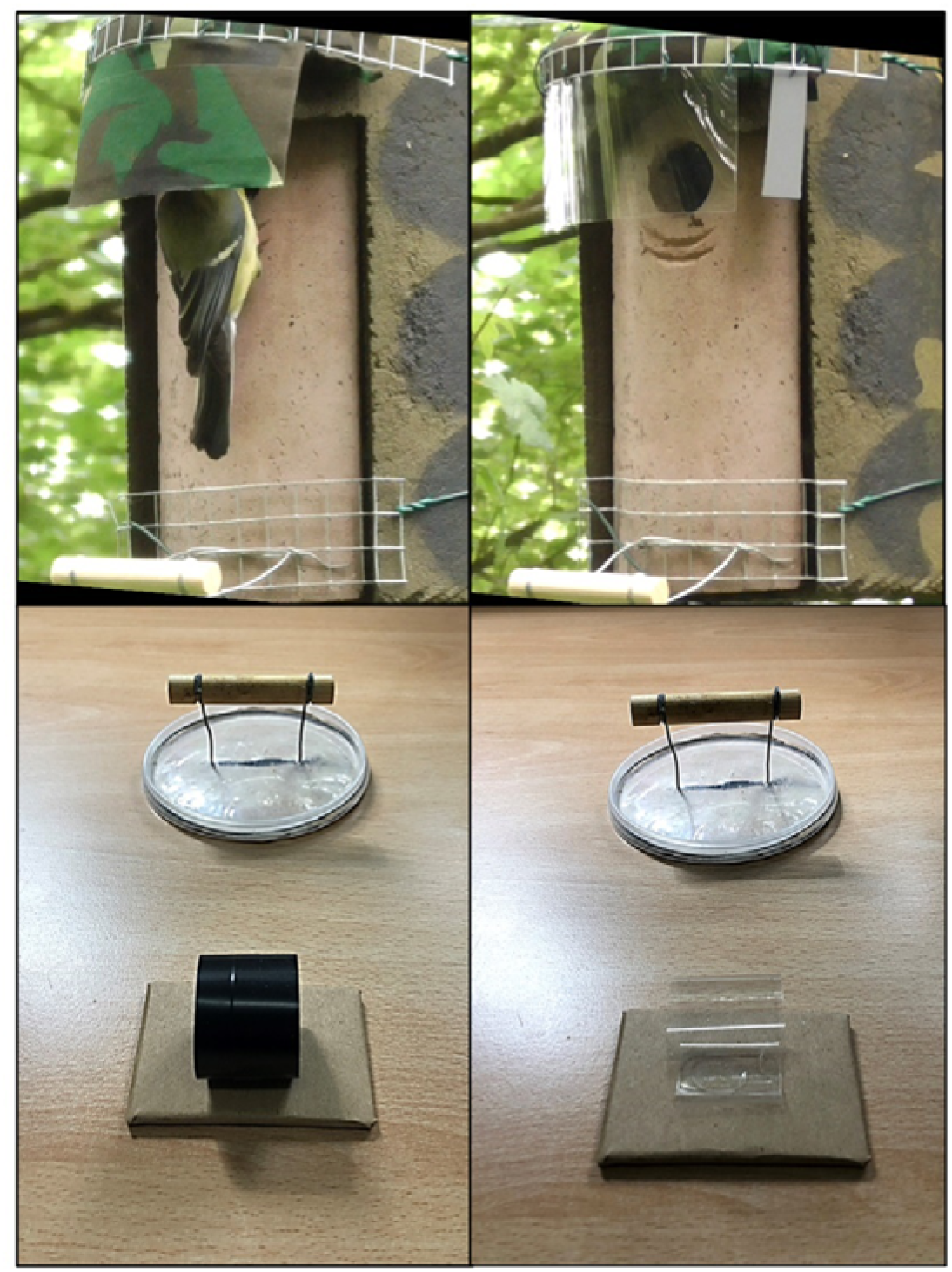
The phases of the detour task in the wild (top) and in captivity (bottom). Training phases with an opaque barrier (left) and test phases with a transparent barrier (right). Perches were positioned below the nest box entrance hole, and in front of the cylinders. For the habituation phase in the wild, the opaque barrier was inverted 180° so it was positioned above the box. For the habituation phase in captivity, the reward was placed visibly on the side of the tube.

At the start of an experimental session (i.e. when the experimental apparatus was placed on the nest box for each phase), the standard nest box front was replaced with an identical-looking front with integrated infrared and RFID technology that logged visits from birds with or without PIT tags (RFID logger for Schwegler 1B from Biomotors Inc). A panasonic HC camera was mounted on a 50cm high tripod, positioned approximately 10 metres from the nest. Both the nest front and the camera were synchronised and used to quantify the number of times each individual bird landed on/entered the nest hole, and to identify the parents by colour ring and/or plumage characteristics in cases when their PIT tag was not in proximity to the reader (i.e. when they touched the barrier). Each experimental session was presented at the nest box for approximately one hour each day over consecutive days. Videos and infrared/RFID data were reviewed on the same day to determine whether birds progressed to the next experimental phase (see criterion details below). In year one, all approaches during the training phase were scored from video, and in year two, videos during the training phase were used only to confirm RFID data.

During the habituation phase, birds were familiarised with the components of the experimental apparatus. These consisted of an opaque rectangular piece of PET plastic film cut from a mobile phone screen protector (0.04W x 10L x 5H cm) covered in camouflage tape attached to the top of the box above the nest hole, a wooden perch positioned at the base of the box attached with metal mesh and wire, and an RFID-equipped nest box front (Figure 1). To pass the habituation phase, individuals had to enter the nest box at least 3 times before moving onto the training phase.

During the training phase, the opaque barrier was inverted so that it covered the nest hole (Figure 1). This phase ensured birds could perform and were familiar with the motor action of going under the barrier. Birds were visually observed from the camcorder footage, and if they went under the barrier without touching it a minimum of three times, they were advanced to the test phase the following day. During the test phase, the barrier was transparent (i.e. without camouflage tape on the plastic film) (Figure 1). The test phase was performed on consecutive days until birds had made five attempts to enter the nest hole, or until a maximum of six days of experiments (including all phases). These attempts, hereafter referred to as ‘trials’, were defined as either colliding with or touching the transparent barrier or accessing the nest hole under the barrier without touching it.

We measured the wild detour task performance as the proportion of successful trials out of total trials, where higher values indicated better performance. A bird was successful if they flew under the barrier, by flying lower than normal or by landing on the perch/box and jumping under. A bird failed if they touched the barrier either by flying into it, by jumping into it from the perch, or by perching on the side and tapping the edge with their beak. On some occasions the lip of the underside of the barrier touched the bird’s back as they jumped under, but this was not considered a fail as it was a clear attempt to avoid the barrier, and likely an artefact of the bird’s size rather than a lack of inhibitory control. If a bird was perched at the nest hole and under the barrier, but did not enter the box, they were still scored as having completed a successful trial. Birds had to fly away from the nest box (i.e. out of the camera view) for a new trial to be scored. If a bird repeatedly jumped between the nest hole and the perch (either contacting the barrier or not), their score was based on their first action, which reflects our scoring criteria in the captive task (see below).

If both parents did not pass a phase after two days, the experiment was abandoned. If only one parent passed the phase after two days, then the experiment was advanced for the participating parent. Therefore, the experiment took between three to six experimental sessions (i.e. days) to complete in year one (mean =3.7 days ±0.13 Standard Error). Experiments started at day ten post-hatching, except for twelve boxes in year one that started at day five-seven due to logistical constraints. In year two the experimental protocol was refined to reduce the number of experimental trials by eliminating a need for a habituation phase, and reducing the criterion for the training phase. Instead of the hour-long habituation phase described above, a dummy apparatus was placed on the box permanently from day eight post-hatching until the start of the training phase. This consisted of wire mesh around the top of the box attached to a plastic rectangle covered with camouflage tape (0.2cm x 6cm x 5cm), and a wooden perch below the nest hole. The criterion for passing the training phase was reduced to one successful approach, instead of three, as our year one data showed that the number of training trials did not influence performance during the test phase (see also results). The experiment took between two and four days to complete in year two (mean = 2.4 days ± 0.12 S.E).

In year one, 19 males and 24 females from 29 nests participated in the experiment. The experiment was attempted at three additional nests, but these were excluded because neither parent passed the habituation or training phases. In year two, 21 females and 20 males from 22 nests participated, where at least one parent from all nests reached and participated in the test phase.

### Experimental procedures: Detour task in captivity

The captive detour task consisted of the same three experimental phases described above. We piloted different sizes of tubes on a cohort of birds not included in the main analyses and chose a 3.5 cm D x 3cm W cylinder tube so that the complexity of the task did not cause ceiling or floor effects in performance caused by the difficulty in detouring around the barrier (Farrar et al., 2020; Völter et al., 2018). A 5 cm high perch was positioned 15 cm in front of the cylinder to standardize the approach direction for each trial (Figure 1). The habituation phase was presented the day after birds arrived in the aviary. Birds had to retrieve a wax worm placed at the edge of an opaque plastic cylinder three times before they received the training phase. Wax worms were euthanised by head compression, so they did not move during the trial. Depending on the bird’s progress, the training phase occurred either on the same day, or the following day.

In the training phase, birds had to retrieve a wax worm placed in the centre of the same plastic cylinder without touching the exterior of the cylinder four out of five consecutive attempts to retrieve the worm. The test phase was always performed the day after a bird passed the training phase. During this phase birds received ten trials with a transparent cylinder of the same dimensions as the one in the habituation and training phases. To ensure birds did not become sated, half a wax worm was used as a reward during the test trials. A trial was defined as a bird approaching the cylinder and making contact with the barrier (scored as a fail) or retrieving the worm from the side of the tube without making contact with the barrier (scored as a success). The trial ended when the bird retrieved the worm, or flew away from the apparatus, at which point the tube was removed from the testing enclosure, rebaited by the experimenter, and placed back in the enclosure. This procedure was designed such that each approach was measured as a success/fail, as opposed to number of pecks until success, the latter of which may be guided by individual persistence (Van Horik et al., 2018). Allowing birds to consume the worm whether they failed or succeeded at each trial controlled for reward history that may have influenced reinforcement and/or motivation through hunger. All birds ate between eight and ten worms at the test phase, except two birds (four worms, seven worms). We recorded the duration it took each bird to complete ten trials as there was no limit to how long birds had to approach the cylinder for each trial. 35 birds participated in the experiment. One additional bird did not settle in the cage or approach the apparatus and therefore did not participate in the experiment. As for the wild task, we measured the captive detour task performance as the proportion of successful trials out of total trials, such that high values indicated putatively high inhibitory control.

### Experimental procedures: Exploration Behaviour

The morning following the birds’ arrival to the aviary, we performed an ‘exploration in a novel environment’ assay, henceforth referred to as exploration behaviour (see also Coomes et al., 2020; adapted from Dingemanse et al., 2002). An access hatch at the back of the bird’s cage that led to a larger room (4.60 m W x 3.10 m L x 2.65 m H) was opened. The light in the home cage was turned off and birds were free to enter the room. Once the bird entered the room, the number of hops and flights within and between trees were recorded from the adjacent corridor through one-way glass. ‘Trees’ were made of a wooden upright support and two thick dowels running at right angles to each other (see also Coomes et al., 2020). The trial was complete after 2 minutes, at which point the birds were returned to their home cage. Exploration behaviour was recorded as the sum of the number of hops and flights, and has been shown to be repeatable in our population (O’Shea et al., 2017).

### Statistical analysis

All models, unless otherwise specified, were run as Generalised Linear (Mixed) Models in lme4 (Bates et al., 2014) in the R statistical software interface (R Core Team, 2014). P values were generated using lmerTest (Kuznetsova et al., 2017). Plots were generated using ggplot (Wilkinson, 2011). We used the dredge function from the MuMIn package (Barton 2019) and an information-theoretic approach in combination with model averaging (Grueber et al., 2011). We generated models from a global model from our GLMMs and retained models with an Akaike’s Information Criterion corrected for small sample sizes (AICc) within 2 units of the top model. We report the conditional averaged weighted parameter estimates across the retained models. All continuous variables were scaled. We used the vif function in the usdm package (Naimi et al., 2014) to test for collinearity between fixed factors. All variables had Variance Inflation Factor less than 2.5 and therefore were not considered to show multicollinearity. Our R code is included in supplementary materials.

### Detour task in the wild

The number of trials undertaken by each bird during the hour test phase varied (mean = 9.6 visits, +/- 0.57 se, 1 min, 26 max). We chose to include a maximum of ten trials as this was consistent with the number of trials in the captive task and existing literature (MacLean et al 2014). We also confirmed that the number of trials used to calculate overall performance did not bias wild detour task performance, if, for example, birds with more trials had higher scores if they learned over successive trials to avoid the barrier. We found that the total number of test trials did not correlate with overall performance using Kendall’s tau correlation test from the cor.test function in R (z=-0.23, tau=-0.02, p = 0.79, n = 84). Moreover, for birds that completed at least ten trials, their overall performance calculated from the first five trials was highly correlated with their overall performance calculated from the first ten trials. (z = 9.42, tau = 0.86, p-value <0.001, n = 69). Eight individuals completed less than five trials and were included in the analysis (3 birds completed 1 trial, 1 bird completed 2 trials, 1 bird completed 3 trials, and 3 birds completed 4 trials).

Initially we examined what fixed effects had a potentially confounding influence on detour task performance in the wild. Lack of any strong effects would lend support for the wild detour task being a reliable test of inhibitory control. It is also important to identify which fixed effects could be driving between individual variation in detour task performance (see repeatability below). We modelled wild detour task performance in a GLMM with a binomial distribution and logit link function, with the number of successes as the numerator and the total trials as a denominator (in R, using cbind, the response variable is entered as two variables, number of successes and number of fails). Our global model included the following fixed effects as potential sources of variation in performance: the number of training trials because the motor action of flying under a barrier could carry-over to the test phase; wing length because size and agility may influence the ability to fly under the barrier; year, lay date and brood size as these may be sources of motivation that may influence parental impulses to feed their chicks; and sex, which has previously been reported to predict inhibitory control performance in a stop-signal task (Lacreuse et al., 2016). Continuous variables were scaled and mean-centred to zero. Site, nest and bird identity (ID) were included as nested random terms. Due to convergence issues associated with overfitting the model with categorical variables, we did not include habitat (conifer versus deciduous) or age in our global model, though visual inspection of these variables and reduced models in which these variables were included suggest they had no effect on performance (Figure S2a, supplementary).

### Detour task in captivity

We modelled captive detour task performance in a GLMM with a binomial distribution and logit link function. Our global model included reward history (i.e. number of worms eaten), motivation (i.e. time to complete all ten test trials), habitat, personality, sex and age as fixed factors, and site of capture as a random effect. Habitat was included as a potential ecological confound linked to differences in population density (O’Shea et al., 2018) and/or environmental variability (van Horik et al., 2019). Exploration behaviour was included as a fixed factor as personality may influence how birds engage with the task, or form part of so-called cognitive ‘styles’ (Sih & Del Giudice, 2012). Sex and age were also included as sources of variation in task performance (Lacreuse et al., 2016; Macdonald et al., 2014). We also investigated whether persistence and captive detour task performance were correlated. To obtain a measure of persistence, we measured the number of times birds pecked at a small transparent case containing a visible but inaccessible mealworm (*Tenebrio molitor*). This data was collected as part of a separate study on foraging choices (Coomes et al., 2020). Persistence data was available for a subset of birds (n=27); therefore, we ran a separate analysis to test for a correlation between the two variables using Kendal’s Tau method for non-normal data with the cor.test function in R.

### Repeatability

We investigated whether individuals showed consistent differences in the wild detour task performance across years (temporal repeatability; but not for the captive task for which we had no repeats). Significant temporal repeatability in performance would suggest that the task measured an intrinsic trait, indicating a permanent environment effect and/or heritability (e.g. Quinn et al., 2009). Our dataset included repeat measures (n=16 observations, 8 individuals), as well as single measures (n= 68) to increase power (Martin et al., 2011). We ran a GLMM as described above, with year as a fixed effect and bird ID as a random effect, and compared this model with another that excluded bird ID using the anova function in R. The repeatability estimate was calculated from the variance components and the residual variance as 1/p(1-p), where p is the expected probability of success calculated as the mean wild detour task performance in the dataset (Nakagawa et al., 2017). 2.5% and 97.5% Confidence Intervals (CI) were calculated with the function confint() using the bootstrap argument with 1000 simulations. Sigma (residual deviation) was estimated to be 1. We also tested whether the inclusion of fixed effects resulted in any change in the repeatability estimate (adjusted repeatability). If the repeatability of wild detour task performance remained significant after inclusion of these effects, this would point to consistent individual differences being explained by inhibitory control.

We also estimated contextual repeatability between tasks, significant levels of which would suggest that performance on these tasks could be attributed to a common factor, supporting the hypothesis that these tasks reflect, at least in part, inhibitory control where the prepotent impulse to go directly towards a positive stimulus (either a food reward or a begging offspring), must be inhibited. We ran an additional two GLMMs (with and without bird ID as a random effect) using a dataset that included repeat measures (n = 21 observations, 10 individuals, one of whom was measured both in year 1 and year 2 of the wild task), and singletons (n=98). We included task (wild versus captivity) as a fixed term, and bird ID and site as random terms. We then repeated these analyses to control for fixed effects that were common between tasks and had been retained in the model selection for temporal repeatability, to ensure that any significant repeatability was not driven solely by common factors unrelated to inhibitory control.

Research and Animal Ethics. This study was conducted under licences from the Health Products Regulatory Authority (AE19130_P017), The National Parks and Wildlife Services (C11/2017, 004/2017, C02/2018 and 001/2018) and the British Trust for Ornithology. The research project received ethical approval from the Animal Welfare Body at University College Cork, and was in accordance with the Guidelines for the Treatment of Animals in Behavioural Research and Teaching (2020).

## Results

### Detour task in the wild

The number of times individuals successfully went under the opaque barrier during the training phase varied across individuals (mean = 7.11±0.59, min 1, max 37) showing that they passed this stage of the task. In year one, during the training phase there were 8 instances from 7 individuals when birds made contact with the opaque barrier, whereas during the test phase there were 130 instances from 36 individuals when birds made contact with the transparent barrier (mean = 3.02±0.45 SE), confirming that the transparent barrier evoked the prepotent response of flying straight to the nest hole (cf. Van Horik et al., 2018). The mean±SE wild detour task performance was 0.56±0.05 SE in year one, and 0.62±0.04 in year two, suggesting no difference between years. Lay date, sex and wing length were retained in the top models, but evidence that any of these terms had an effect on detour task performance was weak because none of them were statistically significant (Figure 2, Table 1), although males performed marginally worse than females, and there was a tendency for performance to decline with lay date. The number of training trials, brood size and year were not retained in any of the top models, and did not predict task performance (Figure 2, Global model test statistics are provided in Supplementary Table 2).

**Figure 2.**
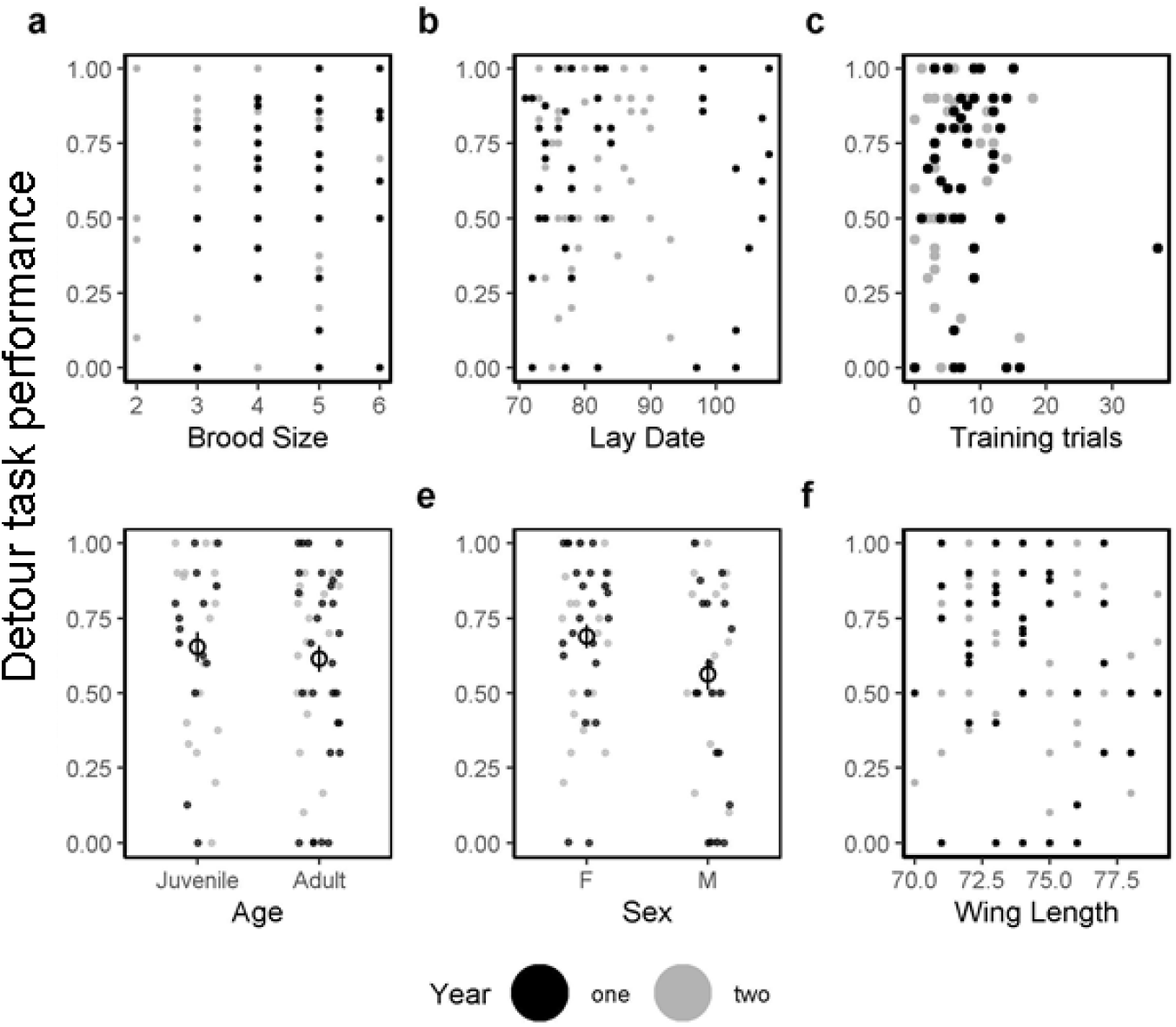
Detour performance (the proportion of successful trials out of total trials, where higher values indicate better performance) in the wild for (a) brood size, (b) lay date (days since 1^st^ March), (c) total number of training trials, (d) age, (e) sex (F=female, M=male), (f) wing length (mm). Points represent individuals where black points were taken in year one, and grey in year two. Datapoints for c, d and f are jittered along the horizontal axis to reduce overlap. Open circle and line represent mean ± standard error across both years. Lay date and sex tended towards significance, and wing length was retained in the final model (Table1).

**Table 1.** Binomial GLMMs of detour task performance (proportion of successful test trials) for the wild detour task (n=84) and the captive detour task (n=35). The values shown are the conditional average of the top models within two AICc of the best model. All continuous variables were scaled. Reference categories for binary variables are in brackets.

| | Estimate ± Standard Error | z | p value |
| --- | --- | --- | --- |
| **Wild detour task** |  |  |  |
| Intercept | 0.96±0.38 | 2.54 | 0.01 |
| Lay date | -0.34±0.20 | -1.65 | 0.10 |
| Sex (Female) | -0.867±0.49 | -1.73 | 0.08 |
| Wing length | 0.35±0.27 | 1.27 | 0.20 |
| **Captive detour task** |  |  |  |
| Intercept | -0.32±0.20 | -1.57 | 0.11 |
| Time to complete test | 0.14±0.12 | 1.23 | 0.25 |
| Sex (Female) | -0.24±0.24 | -0.96 | 0.34 |
| Exploration behaviour | 0.09±0.13 | 0.66 | 0.51 |
| Total rewards eaten | 0.19±0.13 | 1.36 | 0.17 |

### Detour task in captivity

The mean±SE captive detour task performance was 0.41±0.04 SE, substantially lower than that observed in the wild task. The time it took birds to complete the test phase, the number of worms eaten, sex and exploration behaviour were the retained fixed terms in the top models, but evidence that any of these variables had an effect was weak because none were statistically significant (Figure S2). Age (Figure S2d) and habitat (Figure S3b) were not retained in any of the top models and did not predict task performance (Global model test statistics are provided in Supplementary Table 2). There was no relationship between captive detour task performance and our independent measure of persistence, the number of pecks birds made to an inaccessible worm in a transparent casing (z=- 1.06; tau = −0.15; p = 0.29, n = 27).

### Repeatability

The temporal repeatability of wild detour task performance across years was low but significant (R= 0.19, p <0.001, CI =0.06, 0.22), and remained significant when controlling for sex, lay date and wing length (adjusted R = 0.15, p < 0.001, CI= 0.001, 0.18) (Figure 3a, c). Contextual repeatability in performance across tasks was low but significant (R = 0.18, p <0.001, CI = 0.09, 0.24), and remained significant when controlling for sex (adjusted R= 0.17, p <0.001, CI = 0.001, 0.22) (Figure 3b, d).

**Figure 3.**
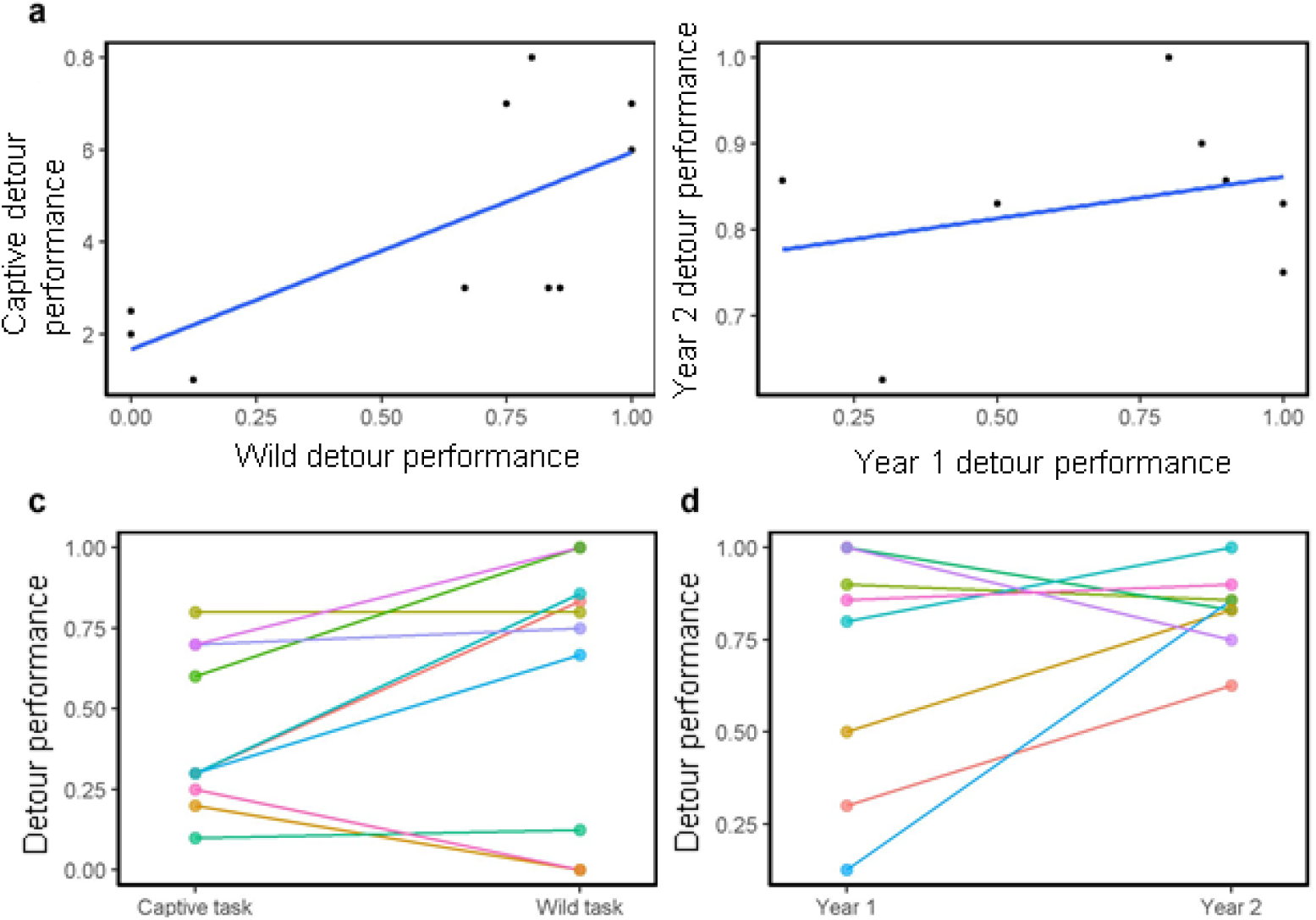
The relationship between individual detour task performance (the proportion of successful trials out of total trials, where higher values indicate better performance) between (a) the wild and captive detour tasks, and (b) year 1 and year 2 wild detour tasks. Reaction norms for (c) contextual repeatability across captive and wild detour tasks, and (d) temporal repeatability across years with the wild detour task. Circles represent individual performance, a regression line is fitted in (a) and (b), and lines (c) and (d) connect repeated measures within individuals in (c) and (d). Lines and circles are in different colours per individual. Repeatability was significant for both analyses, indicating that rank order differences across tasks are more similar than expected by chance.

## Discussion

We show that the detour task is repeatable across years and testing environments (wild versus captivity). Furthermore, we controlled for a range of possible confounding variables, and they did not explain the repeatability of performance. To our knowledge, our study is one of two other known studies that have investigated repeatability of inhibitory control in wild birds (Ashton et al., 2018; Shaw, 2017a), only one of which also reported significant repeatability (Ashton et al., 2018). Moreover, our findings contrast with recent reports that the detour task is confounded by cognitive and/or non-cognitive traits unrelated to inhibitory control (Shaw, 2017b; van Horik et al., 2020; Van Horik et al., 2018). Instead, our study lends support for the detour task as a robust measure of inhibitory control in great tits, at least in the context of inhibiting a motor action. By synthesising our findings across both wild and captive versions of the detour task, we discuss the extent to which we can attribute task performance to inhibitory control.

Performance on the wild detour task was repeatable across time, and between different tasks. Although repeatability estimates were low, these results point to an underlying, inherent trait that was consistently measured despite differences in year, season, testing location and apparatus-type. Repeatability sets the upper limit for heritability but does not preclude permanent environment effects driving some or even all the intrinsic differences observed. Nevertheless, genetic pedigree studies have shown that motor inhibition is heritable in birds (Langley et al., 2020), and that executive functioning is the most heritable psychological trait in humans (Friedman et al., 2008). While we acknowledge that the relatively small sample size of within-individual measurements may render parameter estimation less reliable (Nakagawa & Schielzeth, 2010), our repeatability estimates were consistent with previous reports in animal cognition. A meta-analysis of repeatability estimates across a range of different cognitive tasks including inhibitory control, problem solving, discrimination and reversal learning, memory, physical and spatial cognition found low to moderate R values for temporal repeatability (0.15 and 0.28) and contextual repeatability (0.20-0.28) (Cauchoix et al., 2018). The two known detour task studies in the meta-analysis had very opposing results: no repeatability in New Zealand Robins (*Petroica australis*) (R = 0.002) (Shaw, 2017b) but high repeatability in Australian Magpies (*Cracticus tibicen*) (R =0.80) (Ashton et al., 2018). This suggests that repeatability of the detour task across species and populations may be considerably variable if temporary environmental effects vary across space and time. The particularly high repeatability reported for the Australian Magpies, for example, is likely due to the repeats being taken just two weeks apart (in comparison to the 12 months in this study) which is likely to inflate differences among individuals due to transient factors (Bell et al., 2009; Cole et al., 2011). Disparities in performance and effect sizes across studies are common in detour task studies, and comparative cognition studies generally (Farrar et al., 2020). Our finding that the same individuals performed better on the wild compared to the captive task highlights the difficulty of labelling ‘better’ or ‘poorer’ inhibitory control abilities when comparing performance within or between species across variations of the detour task. Nevertheless, the significant repeatability, coupled with the lack of evidence that extraneous factors had any effect on repeatability, suggests an underlying trait that is common across tasks in our system, which we attribute to being the inhibition of a prepotent response/habitual behaviour.

Motivation (Shaw, 2017b), persistence (Van Horik et al., 2018), and experience (van Horik et al., 2019, 2020) have been proposed to interfere with estimating individual differences in inhibitory control. The use of food rewards may contribute to motivational effects attributed to between-individual differences in hunger state, body condition and/or food preferences (Cooke et al., 2021; Shaw, 2017b). Placing the detour task in front of the nest box access hole allowed us to measure performance independently of food rewards. Moreover, performance was repeatable despite different reward incentives between wild and captive tasks. Although we expected that all individuals would be highly motivated to participate in and complete the inhibitory control tasks as quickly as possible, we nevertheless expected that motivation could vary depending on the reproductive value and the viability of the parent’s offspring. However, we found no evidence for this because performance was unaffected by brood size, brood age and lay date. Approach latencies, or time to complete a task, have been interpreted as proxies for motivation to obtain a reward, but we found no effect of these variables on performance in our captive task, nor did a similar study with common pheasants (*Phasianus colchicus*) (van Horik et al., 2020). Body condition may reflect an individual’s energetic state, and has been linked with performance in the detour task in wild New Zealand robins (Shaw, 2017b). We did not test for such an effect in the current study as we did not have an accurate measure of body weight at the time of the experiments. Weights were only taken when birds were handled (i.e. at capture), and can fluctuate as much as 5% for great tits in captivity (personal observation), and while parents are provisioning chicks. Notwithstanding the point that individual differences in motivation could theoretically remain due to intrinsic differences in motivation (Morand-Ferron et al., 2016), even when body condition is the same, future studies could integrate a weighing scale into the perch placed in front of the cylinder or the nest box to obtain real-time weight as a proxy for motivation due to energetic state.

Persistence, in which animals continue to attempt to obtain a reward, despite persistent errors that do not lead to a reward, has been suggested to contribute to performance in problem solving tasks (Griffin et al., 2015) as well as the detour task (Van Horik et al., 2018). Using data that overlaps with that used here, we found no evidence that persistence in an independent foraging task was related to captive detour task performance (Coomes et al., 2020). The lack of an effect of persistence may be because we scored the initial response (success/fail) for each independent approach to the barrier to quantify inhibitory control, as opposed to measuring performance as the number of pecks before moving around the barrier, the latter quantification being more sensitive to individual differences in persistence, and very similar to how we measured persistence. Indeed, whether behaviour is repeatable or correlates with other traits can be dependent on subtle differences in how behaviour is measured generally (Carter et al., 2012; Davidson et al., 2018).

Between-individual differences in animal personality may influence task performance if personality is intrinsically linked to so-called cognitive “styles” (Sih & Del Giudice, 2012). For example, behaviours commonly attributed to the reactive-proactive animal personality axis include slow exploring, environmentally sensitive individuals on one extreme, and fast exploring, routine forming individuals on the other (Réale et al., 2007). These definitions have many parallels with definitions associated with human inhibition and impulsivity, including ‘an impulsive behaviour with no forethought of consequences’ (Moeller et al., 2001). We found no evidence that exploratory behaviour, a common behaviour associated with the fast-slow exploration personality axis, was associated with performance on the detour task in captivity, thus excluding this as an influence on performance in the detour task. This is consistent with reports in black-capped chickadees (*Poecile atricapillus*) (Guillette et al., 2017) and domestic dogs (*Canis lupus familiaris*) (Bray et al., 2015), but not common waxbills (*Estrilda astrild*) (Gomes et al., 2020). We note that our measure of personality is purely an index of the fast-slow personality axis (Bell, 2007), and it may be that specific facets of this axis, for example responsiveness, the quality of exploration (i.e. information gathered) and neophobia, need to be measured in isolation to detect links with inhibitory control. Similarly, other kinds of personality axes, or behavioural variation in general, could play a role. Equally, we also note that our measure of inhibitory control likely only captures one facet of self-regulation but there are many others (e.g. delayed gratification; e.g. Mischel et al., 1989) that themselves may be controlled by distinct but related forms of inhibitory control we measure here. Despite the challenges in teasing apart different elements of inhibitory control, and cognition generally, much of the psychology literature suggests that different measures of inhibitory control are a component of a wider latent cognitive variable, such as general inhibition, executive functioning and general intelligence (Anderson & Weaver, 2009; Aron et al., 2004; Ashton et al., 2018; Bari & Robbins, 2013; Shaw et al., 2015), but it remains to be seen whether that is also true in wild study systems.

In both detour tasks, performance on successive trials could have been driven by motor learning from previous trials. The acquisition of a motor routine has been shown to affect detour task performance in pheasants (van Horik et al., 2020), but this was not the case in our wild task. It is often assumed that a major drawback of cognitive tests in the wild is that it is difficult to control for previous experience, but our results suggest that the wild detour task is not sensitive to differences in the number of training or test trials in which individuals participate. Fine scale local factors could also drive consistency across years - for example approach direction and perception of the transparent barrier associated with nest box orientation or lighting conditions - but many of these are unlikely because birds typically change their nest box every year, and only one of the nest boxes was occupied in both years in our sample. Nevertheless it is highly likely that a range of other confounding effects could have influenced performance in our tasks, some of which are likely to be normally overlooked even in controlled experiments (e.g. Dunn et al., 2011).

The field of animal cognition has made major advances in describing cognitive variation between and within species, yet obtaining unbiased and realistic estimates of cognitive variation in natural populations remains a significant challenge (Morand-Ferron et al., 2016; Rowe & Healy, 2014; Thornton et al., 2014). Limited participation in self-administered cognitive trials in the wild potentially leads to bias towards some kinds of individuals, for example those with higher cognitive abilities (Reichert et al., 2020). By deploying a modified version of the detour task that we developed specifically for our system, where birds were compelled to visit, we were able to minimise participation bias at the population level, which can have a big impact on parameter estimation generally. Additionally, our approach limited the effects of human interference and social interactions by conducting the task at isolated locations. Finally, our results support the traditionally held view that the detour task is a reliable measure of inhibitory control, a cognitive process that is likely an important driver of functionally important behavioural plasticity. While it may never be feasible to study inhibitory control as a discrete module in isolation from extraneous/integrated processes, which is true for many cognitive processes, it may not be advisable or necessary to do so when addressing questions in evolutionary ecology (Morand-Ferron & Quinn, 2015) since selection rarely acts on individual genes. Overall, cognitive estimates derived from the detour task can hold value either as a stand-alone task specifically measuring inhibition, or as part of a larger test battery aimed at understanding general cognitive ability.

## Author contribution

G.L.D. designed the experiment with input from all authors. G.L.D. ran the experiments with assistance for M.S.R., J.R.C., I.G.K and I.D.H. G.L.D analysed the data. G.L.D and J.L.Q wrote the manuscript with input from all authors.

## Acknowledgements

We wish to thank Sam Bayley, Jodie Crane, and James O’Neill for assistance with fieldwork, and Amy Cooke for assistance in the aviary. Thank you to Karen Cogan, Allan Whitaker and Luke Harman for technical support. Funding came from the European Research Council under the European Union’s Horizon 2020 Programme (FP7/2007-2013)/ERC Consolidator Grant “EVOLECOCOG” Project No. 617509, awarded to J.L.Q., and from a Science Foundation Ireland ERC Support Grant 14/ERC/B3118 to J.L.Q.

