## Supplementary Materials for "Inhibitory control performance is repeatable across years and contexts in a wild bird population"

Table S1. List of the 10 study sites in the Bandon Valley area, co. Cork (Ireland), where birds were studied. We show the coordinates in decimal degrees (DD) and the main habitat type (mixed deciduous forest vs. coniferous plantation) for each site. *sites that were included in year two; ǂ a site where birds were not caught for the captive task.

| Study site | Habitat type | Coordinates (DD) |
| --- | --- | --- |
| Ballinphelic | Coniferous | 51.840074, -8.628249 |
| Castlebernard* | Mixed | 51.741821, -8.773443 |
| Piercetown | Coniferous | 51.789310, -8.451774 |
| Dukes Wood* | Mixed | 51.787396, -8.755946 |
| Farran | Coniferous | 51.700448, -8.805728 |
| Garretstown | Coniferous | 51.654770, -8.616293 |
| Inishannon* | Mixed | 51.763048, -8.662415 |
| Kilbrittain* | Mixed | 51.671880, -8.683837 |
| Lissarda | Coniferous | 51.858086, -8.859128 |
| Shipoolǂ | Mixed | 51.737620, -8.631437 |

Table S2. Test statistics from the global binomial GLMMs of detour performance (proportion of successful test trials) for the wild detour task (n=84) and the captive detour task (n=35). All continuous variables were scaled. Reference categories for binary variables are in brackets.

|  | **Estimate ± Standard Error** | **z** | **p value** |
| --- | --- | --- | --- |
| Wild detour task |  |  |  |
| Intercept | 1.1±0.43 | 2.56 | 0.01 |
| Brood size | -0.02±0.19 | -0.09 | 0.93 |
| Total training trials | -0.05±0.17 | -0.3 | 0.76 |
| Year | 0.23±0.43 | 0.53 | 0.59 |
| Wing length | 0.36±0.27 | 1.33 | 0.18 |
| Lay date | -0.35±0.2 | -1.77 | 0.08 |
| Sex | -1.19±0.52 | -2.31 | 0.02 |
| Captive detour task |  |  |  |
| Intercept | -0.34±0.27 | -1.3 | 0.19 |
| Sex | -0.18±0.26 | -0.68 | 0.5 |
| Age | 0.16±0.28 | 0.57 | 0.57 |
| Time to complete test | 0.11±0.12 | 0.94 | 0.35 |
| Total rewards eaten | 0.18±0.13 | 1.36 | 0.17 |
| Exploration behaviour | 0.17±0.14 | 1.25 | 0.21 |
| Habitat | 0.11±0.36 | 0.312 | 0.76 |

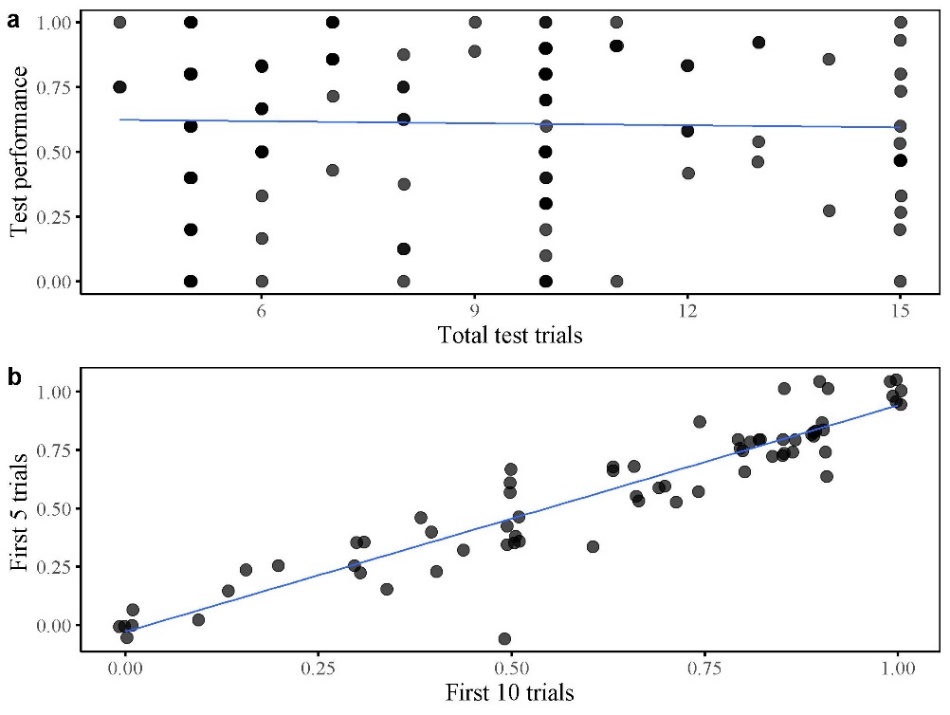

Figure S1. The number of test trials and detour task performance in the wild (the proportion of successful trials out of total trials, where higher values indicated better performance). Circles represent individual’s overall performance where a) The number of test trials did not predict performance on the wild detour task, and b) Individual’s overall performance calculated from the first 10 trials was highly correlated with their overall performance if calculated from the first 5 trials.

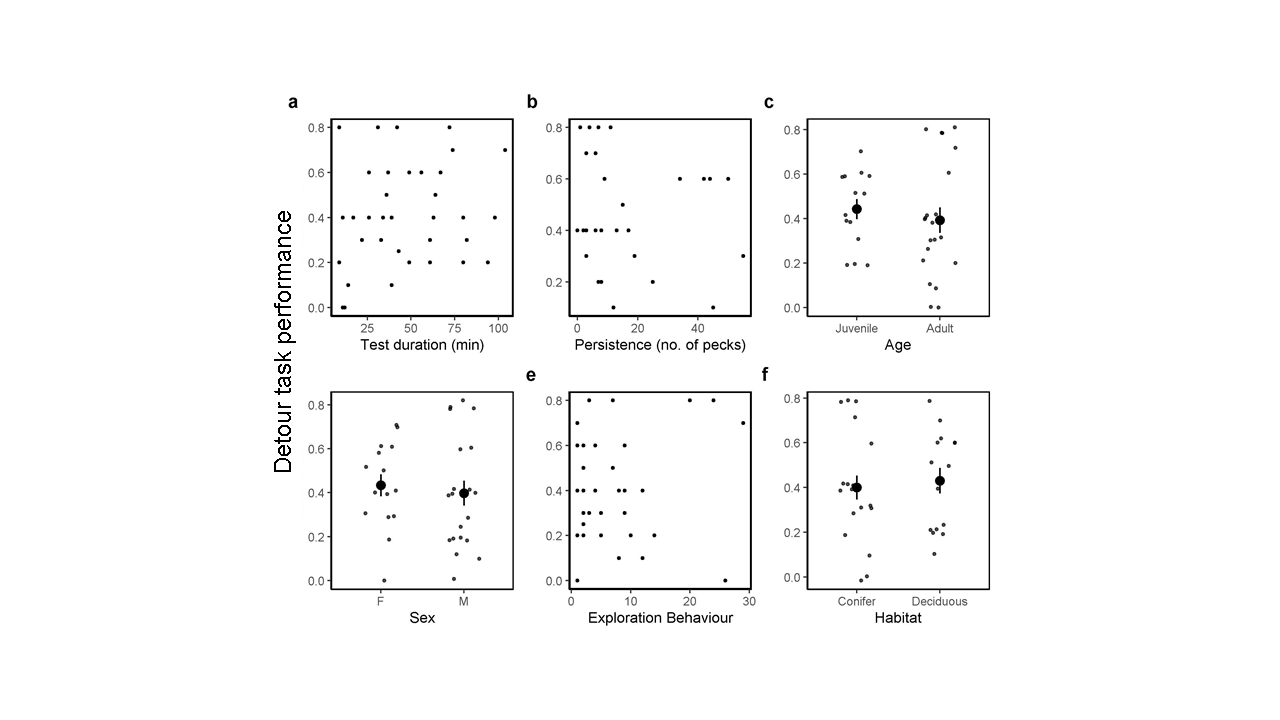

Figure S2. Detour performance (the proportion of successful trials out of total trials, where higher values indicate better performance ) in captivity (a) test duration, (b) persistence, (c) total rewards eaten, (d) age, (e) sex (F=female, M=male), (f) exploration behaviour. Points represent individuals and are jittered along the horizontal axis to reduce overlap in c, d and f. Open circle and line represent mean ± standard error. Test duration, total rewards eaten, sex and exploration were retained in the final model, but were non-significant (Table1)

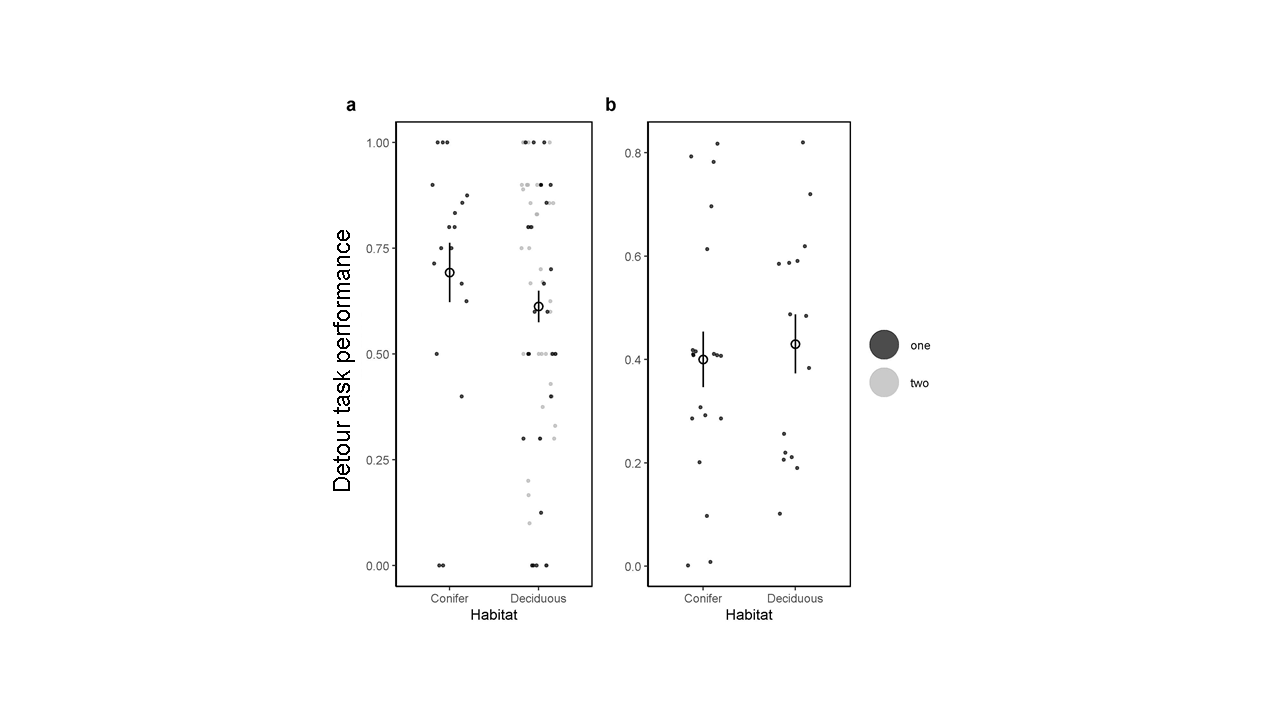

Figure S3. Detour performance (the proportion of successful trials out of total trials, where higher values indicate better performance) across habitats for birds measured in (a) the wild, (b) captivity. Points represent individuals where black points were taken in year one, and grey in year two. Note that no individuals were measured in conifer habitats in year two. Datapoints are jittered along the horizontal axis to reduce overlap. Open circle and line represent mean ± standard error across both years.
